# Mechanism of Arrhythmogenesis Driven by Early After Depolarizations in Cardiac Tissue

**DOI:** 10.1101/2024.11.14.623585

**Authors:** Jack Stein, D’Artagnan Greene, Flavio Fenton, Yohannes Shiferaw

## Abstract

Early-after depolarizations (EADs) are changes in the action potential plateau that can lead to cardiac arrhythmia. At the cellular level, these oscillations are irregular and change from beat to beat due to the sensitivity of voltage repolarization to subcellular stochastic processes. However, the behavior of EADs in tissue, where cells are strongly coupled by gap junctions, is less understood. In this study, we develop a computational model of EADs caused by a reduction in the rate of calcium-induced inactivation of the L-type calcium channel. We find that, as inactivation decreases EADs occur with durations varying randomly from beat to beat. In cardiac tissue, however, gap junction coupling between cells dampens these fluctuations, and it is unclear what dictates the formation of EADs. In this study we show that EADs in cardiac tissue can be modeled by the deterministic limit of a stochastic single-cell model. Analysis of this deterministic model reveals that EADs emerge in tissue after an abrupt transition to alternans, where large populations of cells suddenly synchronize, causing EADs on every other beat. We analyze this transition and show that it is due to a discontinuous bifurcation that leads to a large change in the action potential duration in response to very small changes in pacing rate. We further demonstrate that this transition is highly arrhythmogenic, as the sudden onset of EADs in cardiac tissue promotes conduction block and reentry. Our results highlight the importance of EAD alternans in arrhythmogenesis and suggests that ectopic beats are not required.

## Introduction

Spatial heterogeneities in the action potential duration (APD) are dangerous because they promote wave break. This occurs when an action potential (AP) wavefront fails to propagate in regions of cardiac tissue that have not fully recovered from a previous excitation. When wave break occurs in a localized area, it can lead to reentry and fibrillation(1). Additionally, heterogeneities in action potentials can result in after-depolarizations, which may cause premature ventricular contractions (PVCs) which can propagate and induce reentry(2). Also, early-after depolarizations (EADs) (3-5) are a key contributor to APD heterogeneity. EADs are unusually prolonged action potentials caused by the failure of repolarizing currents to overcome inward currents that prolong the action potential (AP). They typically occur during phase 2 of the AP and can be caused by an increased L-type calcium current (LCC) or a decrease in potassium currents that control cell repolarization. EADs are particularly dangerous because they occur randomly in populations of cells, thereby driving APD heterogeneities in cardiac tissue(6-8). Thus, EADs are often associated with arrhythmias such as Torsade de Pointes, long QT syndromes and heart failure(9-11).

Several studies have investigated the underlying mechanisms of EADs in cardiac cells(12). In particular, Tran et al. (13) showed that EADs are due to oscillations caused by a Hopf bifurcation that occurs during the AP plateau. Also, Bertran et al. (14) applied canard theory to show that EADs occur when the Ca current is enlarged and is caused by the LCC window current. More recently, Wang et al. (15) identified four distinct mechanisms for EADs, driven by voltage dynamics, calcium dynamics, or their interactions. Also, Huang et al. (16) conducted an extensive study on the temporal characteristics of EADs, demonstrating that they exhibit strong beat-to-beat fluctuations. This is because the membrane voltage becomes extremely sensitive to ion currents that maintain the AP plateau, amplifying the inherent stochasticity present at the subcellular level. In cardiac tissue, this stochasticity is substantially dampened due to the electrical coupling between cells via gap junction channels, which effectively average the voltage over large populations of cells in tissue. However, it is not well understood how EADs in tissue differ from those observed at the single-cell level. In an insightful study, Sato et al. (17) demonstrated that EADs in tissue are highly arrhythmogenic, inducing stark APD heterogeneities with complex spatiotemporal dynamics. In a subsequent study, Sato et al. (18) analyzed the sources of these heterogeneities and found that they arise from a combination of inherent cellular stochasticity and dynamical chaos due to the system’s nonlinearity. These findings indicate that EADs in tissue are both highly nonlinear and stochastic, and both factors may play a key role in the formation of cardiac arrhythmias.

In this study, we use a phenomenological model of a ventricular cell to investigate the properties of EADs in both isolated cells and electrically coupled tissue. The model captures subcellular calcium (Ca) release by tracking the stochastic recruitment and termination of Ca sparks. This stochastic behavior is linked to Ca-sensitive membrane channels, resulting in beat-to-beat variability in the APD. Our findings suggest that, to replicate the experimentally observed APD variations in guinea pig myocytes (19), approximately 4,000 ryanodine receptor (RyR) clusters must be available for Ca spark recruitment. Using this model, we study the formation of EADs in electrically coupled tissue and find that APD fluctuations are significantly dampened compared to isolated cells. In this context, EADs are well described by the deterministic limit of the subcellular stochastic model. Our analysis of this model shows that EADs in cardiac tissue manifest as APD alternans, where the long APD corresponds to an AP with an EAD followed by a short AP with no EAD. A detailed examination of this deterministic model reveals that the transition to EAD alternans occurs via a subcritical pitchfork bifurcation. This transition is discontinuous, so that infinitesimal changes in pacing frequency can lead to large changes in the APD response. Moreover, we show that the presence of EADs in cardiac tissue makes the system highly susceptible to conduction block and reentry. Thus, our results suggest that EAD alternans is a key mechanism for arrhythmogenesis, and ectopic beats are not necessary for the initiation of arrhythmic events.

## Methods

### Modeling Ca spark recruitment in ventricular myocytes

In this study we will develop a phenomenological model of Ca cycling in a ventricular cell which incorporates stochastic dynamics. This model is based on a previously developed model of an atrial cell (20,21) and has been modified for the ventricular cell geometry. In our approach, we model the stochastic recruitment and extinguishing of the population of Ca sparks within the cell. Since Ca sparks occur randomly, the number of Ca sparks in the cell constitute the primary source of noise in our model. In this approach we separate RyR clusters in the cell into two groups: (i) A population of RyR clusters, referred to as junctional clusters (J), which are in close proximity to LCC channels on the cell membrane. (ii) Non-junctional RyR clusters (NJ) in the cell interior which are far from LCC channels. Under normal conditions Ca release in the cell is due to Ca sparks recruited at J junctions that are triggered by the nearby LCC channel openings. In contrast, NJ junctions are far from LCCs and do not respond to LCC openings but can release Ca if the local Ca concentration rises independently of LCC openings. The distinction between J and NJ clusters is crucial, as the population of RyR clusters activated by LCCs is typically smaller than the total number of clusters in the cell. This distinction is relevant in this study since the fluctuations in Ca spark recruitment depends on the number of clusters that can be activated by LCC openings.

In ventricular myocytes J clusters are distributed uniformly in the cell since t-tubules penetrate into the cell volume and distribute LCCs inside the cell interior(22,23). In contrast, in atrial myocytes J clusters are distributed mostly on the cell periphery, and Ca release in the interior can only occur due to propagating Ca waves which are due to Ca sparks at NJ clusters(24-26). In this study we will not consider the effect of wave propagation, although this effect can be added to our computational framework. To proceed, we will denote *n*_*b*_(*t*) as the number of J clusters at which Ca is being released due to a Ca spark at time *t*. Following our previous work(20) we model the change in the number of sparks according to the reaction scheme

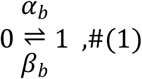

where *α*_*b*_ is the rate of Ca spark recruitment at J sites, and where *β*_*b*_ is the rate at which these sparks are extinguished. During a small time increment Δ*t* the number of Ca sparks in the cell evolves according to

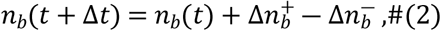

where

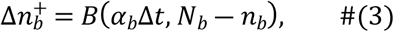

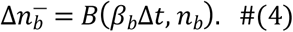

Here, *N*_*b*_ denotes the total number of J clusters, and *B*(*p,n*) is a random number picked from a binomial distribution with success probability *p* and number of trials *n*.

### Cell compartmentalization and Ca cycling

To model Ca dynamics in the cell, we constructed phenomenological equations representing the average Ca concentrations near J and NJ clusters. To develop these equations, we first partition the cell interior into 4 regions which are illustrated in Figure 1A. These volumes are: (i) The total cytosolic volume in the vicinity of J clusters with average free Ca concentration *c*_*b*_ and total volume *v*_*b*_. (ii) The total cytosolic volume in the vicinity of NJ clusters with average free Ca concentration *c*_*i*_ and total volume *v*_*i*_. (iii) The total SR volume in the vicinity of J clusters with an average concentration of *c*_*srb*_ and volume *v*_*srb*_. (iv) The total SR volume in the vicinity of NJ clusters with an average concentration of *c*_*sri*_ and volume *v*_*sri*_. Once the number of Ca sparks is computed using equations 2-4 we set the current flux from the SR into the cytosol as

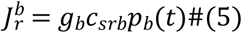

where *p*_*b*_(*t*) = *n*_*b*_(*t*)/*N*_*b*_ is the fraction of J junctions at which a spark occurs, and where *g*_*b*_ is the effective conductance of RyR clusters in the cell. The other Ca fluxes linking these compartments are illustrated in Figure 1B, and the fluxes in Figure 1A are defined in Table 1. All details of our model construction are given in the Online Supplement.

**Figure 1.**
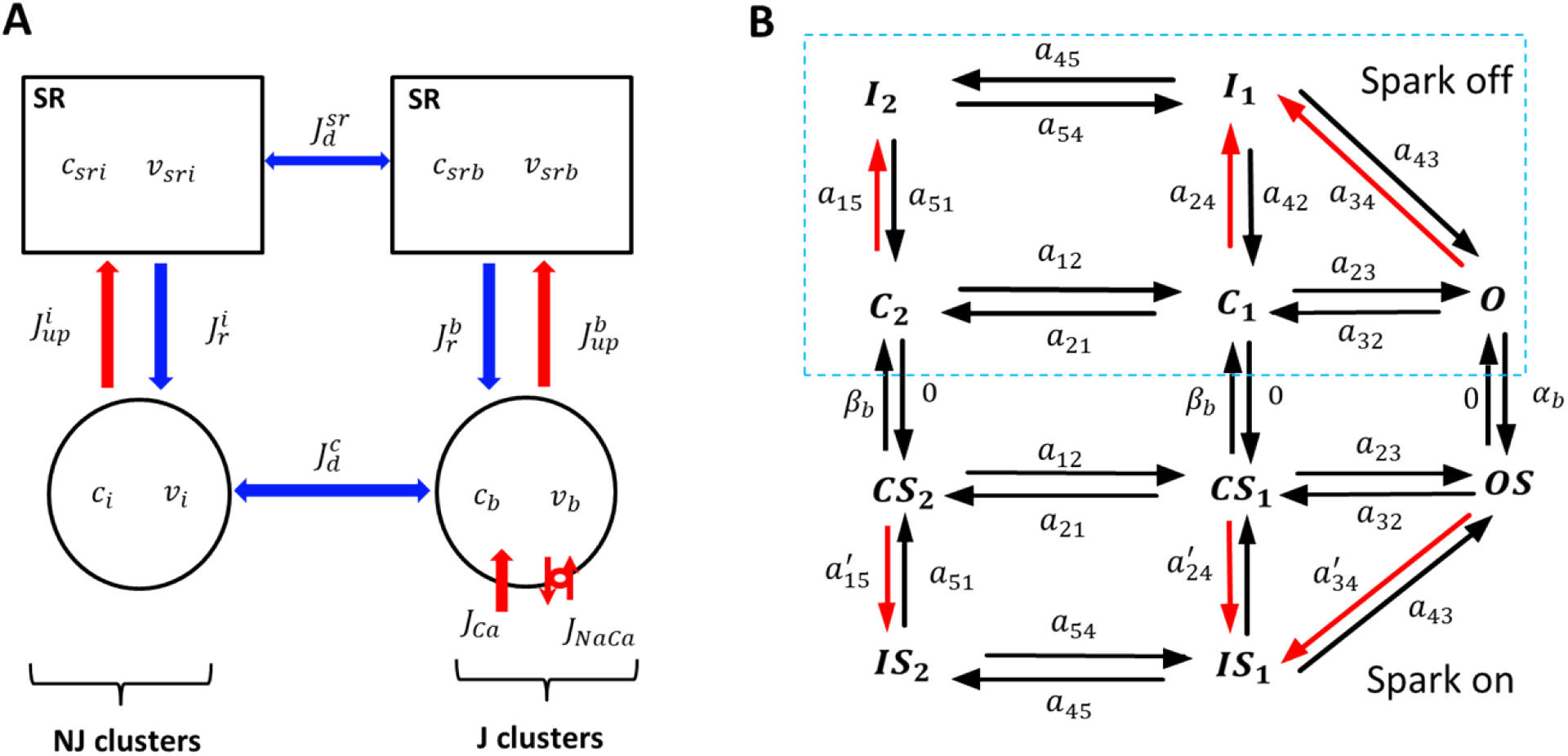
Phenomenological model of Ca spark recruitment in ventricular myocytes. **(A)** Compartment architecture. The cell is divided into 4 compartments which represent the average Ca concentration in the SR and cytosol in the vicinity of J and NJ clusters. The concentrations *c*_*b*_ and *c*_*srb*_ denote the average concentration in the cytosol and SR in the vicinity of J clusters, while *c*_*i*_ and *c*_*sri*_ denote the concentrations near NJ clusters. The total SR volume near J and NJ clusters is denoted as *v*_*srb*_ and *v*_*sri*_ respectively, while the total cytosolic volume is denoted as *v*_*b*_ and *v*_*i*_. The currents linking these two spaces are the Ca release 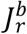 due to a population of sparks at J clusters, and the uptake pump current 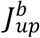 which replenishes the SR volume. Similarly, Ca release and uptake from the SR volume near NJ clusters is denoted by 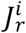 and 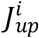 respectively. The currents 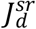 and 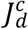 are averaged diffusive currents proportional to the concentration difference between the internal cell volumes. **(B)** Markovian model of the LCC current. Markov states in dashed box (spark off) represent channels facing low Ca (0.1*μM*), while channels outside box (spark on) face a Ca spark with high local Ca (100*μM*). The terms *a*_*ij*_ denote the transition rates between the corresponding states. Red (black) arrows indicate Ca (voltage) dependent transition rates.

**Table 1:** Description of Ca fluxes in phenomenological model.

| Flux | Description |
| --- | --- |
| $J_r^b$ | Total RyR flux from J clusters to the cytosolic space in the cell interior. |
| $J_r^i$ | Total RyR flux from NJ clusters to cytosolic space in the cell interior. |
| $J_{up}^b$ | Total uptake flux from the cytosol near J sites into the SR. |
| $J_{up}^i$ | Total uptake flux from cytosol near NJ sites to the SR. |
| $J_d^{sr}$ | Total diffusive flux from SR volumes near J sites to NJ sites. |
| $J_d^c$ | Total diffusive flux from cytosol near J sites to NJ sites. |
| $J_{Ca}$ | Total LCC flux into the cell. |
| $J_{NaCa}$ | Total Sodium-Calcium exchanger current into the cell. |

### The L-type Ca current

To model the LCC current we will apply a Markovian model, due to Mahajan et al (27), that is based on Rabbit ventricular data. The Markov state diagram introduced in that study, and further developed for population based models(20), is shown in Figure 1B. Briefly, we have split our population of LCCs into two groups of Markov states. The first group, which we will refer to as the “spark off” group, is shown inside the dashed box (Figure 1B) and represents LCCs that are far from active sparks. The Markov states describing these channels consist of 2 closed states (*C*_1_,*C*_2_), one open state (*O*), and 2 inactive states (*I*_1_, *I*_2_), with red arrows denoting Ca dependent rates. Since these channels are not in the vicinity of a Ca spark their Ca dependent transition rates depend only on the diastolic Ca concentration in the cell. On the other hand, LCCs that are in the same junctional space of active sparks (spark on) are described by Markov states (*CS*_1_,*CS*_2_,*OS, IS*_1_,*IS*_2_) which are regulated by a local Ca concentration that is substantially larger. These channels undergo much faster Ca induced inactivation. The transition rates between groups of channels are then modelled by letting open LCCs in the “spark off” group transition to the “spark on” group at a rate that is the spark recruitment rate *α*_*b*_. Similarly, the reverse transition will be set to the rate that sparks extinguish *β*_*b*_. Detailed model parameters describing the channel transitions rates are given in the Online Supplement.

The transition rates in the LCC model are derived from experimental data established in the Mahajan model. This model was constructed by fitting the LCC current using both barium (Ba) and calcium (Ca) as charge carriers. This approach separated the effect of Ca and voltage dependent inactivation, which is controlled by the transition rates *a*_24_ and *a*_34_ (Figure 1B) for LCCs with no local spark, and 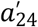 and 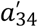 for channels in the vicinity of a Ca spark. These rates are taken to have the phenomenological form

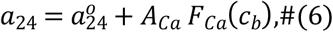

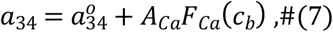

where 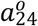 and 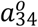 determine the rate of inactivation that is independent of Ca. The Ca dependence of inactivation is modelled as a function of the concentration near J clusters, and is given by

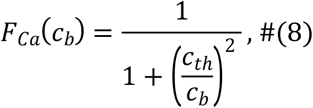

where *c*_*th*_ is the threshold for Ca induced inactivation. Also, the parameter *A*_*Ca*_ controls the relative strength of the Ca dependent and Ca independent inactivation rates. In our approach 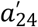 and 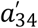 are taken to have the same form, the main difference being that the Ca concentration *c*_*b*_ in the vicinity of a Ca spark is large so that *c*_*b*_ ≫ *c*_*th*_ and we set *F*_*Ca*_(*c*_*b*_) = 1.

### Modeling EADs

EADs have been extensively studied both computationally and experimentally, and it is well established that they can be induced by increasing the LCC (6). Experimentally, this can be achieved by using a calcium channel agonist such as Bay K. EADs are also observed in congenital long QT (LQT) syndrome (9,28,29) and, in some cases, have been linked to mutations in CaM(28,30), which mediates calcium-induced inactivation. To induce EADs in our model we have reduced the component of calcium-induced inactivation of the LCC current by lowering the parameter *A*_*Ca*_ in equations 6-7. This change prolongs the AP and induces EADs at pacing cycle lengths (*CL*) of *CL* ∼ 500*ms*.

### Action potential model

To model cardiac tissue we have integrated our Ca cycling equations with the major ion currents from the Mahajan AP model for the rabbit ventricular myocyte (27). In particular we incorporate their ion current formulations for the fast sodium current (*I*_*Na*_), the rapidly activating delayed rectifier *K*^+^ current (*I*_*Kr*_), the slowly activating delayed rectifier *K*^+^ current (*I*_*Ks*_), the inward rectifier *K*^+^ current (*I*_*K*1_), the transient outward *K*^+^ current (*I*_*to*_), the *Na*^+^/*K*^+^ exchange current (*I*_*NaK*_), and finally the sodium-calcium exchanger current (*I*_*NaCa*_).

### Simulations of 2D cardiac tissue

In this study we will explore the dynamics of electrical propagation in a 2D tissue of cells described by our phenomenological model. To model electrical propagation, we apply the cable equation

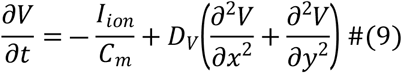

where *C*_*m*_ = 1*μF*/*cm*^2^ is the membrane capacitance, *D*_*V*_ = 1 × 10^―4^*cm*^2^/*ms* is the effective voltage diffusion coefficient, and where *I*_*ion*_ is the total transmembrane current. The cable equation is integrated using an operator splitting approach(31), with a space step Δ*x* = 0.015*cm* and with a variable time step in the range *dt* = 0.01 ― 0.1*ms*.

## Results

### Single cell model calibration and fluctuations of the APD

Our ventricular cell model exhibits fluctuations in the voltage time course, which are due to the stochastic recruitment and extinguishing of Ca sparks. The coupling between subcellular stochasticity and the voltage across the cell membrane is mainly driven by the Ca dependence of the sodium-calcium exchange current (*I*_*NaCa*_). Since Ca sparks are recruited only from J clusters the beat-to-beat fluctuations in the total number of sparks recruited will depend on the number of available clusters *N*_*b*_. To determine *N*_*b*_ we will rely on the experimental work of Zamboni et al. (19) who measured the beat-to-beat fluctuations of the APD of isolated guinea pig myocytes paced at a steady state cycle length (*CL*). Specifically, they measured the APD at 90% repolarization (*APD*_90_) and found that the average over 10 beats, after pacing to steady state (2 minutes at *CL* = 500*ms*), was ⟨*APD*_90_⟩ = 342.8*ms* with a standard deviation *σ* = 10.4*ms*. To characterize the magnitude of the fluctuations with respect to the APD they computed the coefficient of variability *c*_*v*_ = *σ*/⟨*APD*⟩. This measurement was made on 132 myocytes and they obtained *c*_*v*_ = 2.3 ± 0.9%. To align our model with these findings we have adjusted our number of J clusters (*N*_*b*_) so that our measured *c*_*v*_ at steady state is within the experimental range. In our computational model we find that *N*_*b*_ ∼ 4000 reproduces the correct order of magnitude of statistical fluctuations observed experimentally. In Figure 2A we show the voltage time course *V*(*t*) using *N*_*b*_ = 4000. The cell model has been paced to steady state for 50 beats at *CL* = 500*ms*, and *V*(*t*) for the final 20 beats is superimposed on the same graph. In Figure 2B we show the histogram of *APD*_90_ using a long simulation of 15,000 beats. In this case we find that the average ⟨*APD*_90_⟩ = 237*ms* with a standard deviation of *σ* = 6.3*ms* and *c*_*v*_ = 2.7%. Here, we note that there are potentially many sources of noise that cause APD fluctuations. Our study suggests that Ca cycling likely contributes a significant component of this noise. However, we mention here that the main findings of this study do not depend on the precise origin of the APD fluctuations. In the discussion section, we will assess the robustness of our main results and demonstrate that they remain valid regardless of the specific source of the underlying noise.

**Figure 2.**
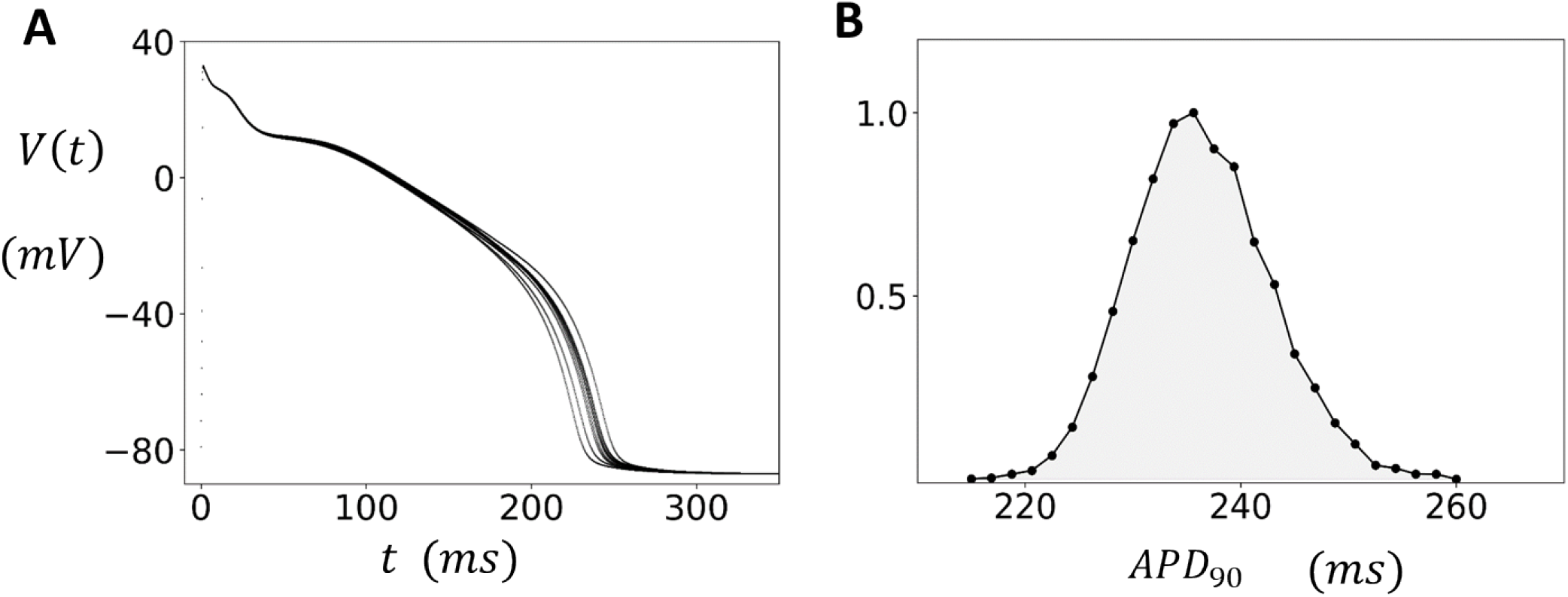
APD fluctuations in the absence of EADs. **(A)** Beat-to-beat voltage fluctuations. The cell is paced at *CL* = 500*ms* to steady state (50 beats), and the voltage of the last ten beats is overlayed on the same graph. The number of J clusters is *N*_*b*_ = 4000. **(B)** Distribution of *APD*_90_. Histogram is constructed by measuring the APD at 90% repolarization over 15000 beats. The distribution is normalized so that the maximum value is 1.

### Statistics of EADs in an isolated cell

In this section, we explore the statistical fluctuations of the APD in the case where the cell model exhibits EADs. To induce EADs we follow the procedure outlined in the methods section, where EADs are induced by lowering the parameter *A*_*Ca*_ in equations 6-7. This reduction increases Ca influx by reducing the component of Ca-induced inactivation, which promotes the development of EADs at longer cycle lengths. When EADs occur, the APD is prolonged and full repolarization does not occur on some beats. Thus, to capture the fluctuations in APD we found it necessary to measure the APD when the voltage crosses a higher threshold of *V*_*c*_ = ―40*mV* during the repolarization phase of the AP. Henceforth, all APDs will be computed using this threshold. In Figure 3A, we plot the measured APD over the last 50 beats after pacing to steady state (200 beats), across a range of *CLs*. The black points represent normal Ca-induced inactivation parameters (*A*_*Ca*_ = 0.15) where EADs do not occur. However, when the component of Ca-induced inactivation is reduced (*A*_*Ca*_ = 0.09) we find that EADs occur at *CLs* larger than *CL*_*c*_ ≈ 500*ms* (red points), so that for *CL* > *CL*_*c*_ EADs occur and large beat-to-beat changes of the APD is observed. In this simulation we have used a dynamic pacing protocol where the *CL* is increased incrementally after each round of 200 paced beats. In Figure 3B, we show an overlay of the voltage time course for the last 50 beats after pacing the cell to steady state at *CL* = 520*ms*. Here, we observe that the repolarization time course is highly variable, with some beats exhibiting EADs characterized by a notch in the AP, while other beats do not. To quantify the distribution of the APD we have measured the APD for 15000 beats at *CL* = 520*ms* and plotted a histogram of the results (Figure 3C). At this cycle length we find that the APD is broadly distributed in the range ∼ 100*ms* ― 500*ms* and is centered around three peaks, which indicates that APD variability increases substantially when EADs occur.

**Figure 3.**
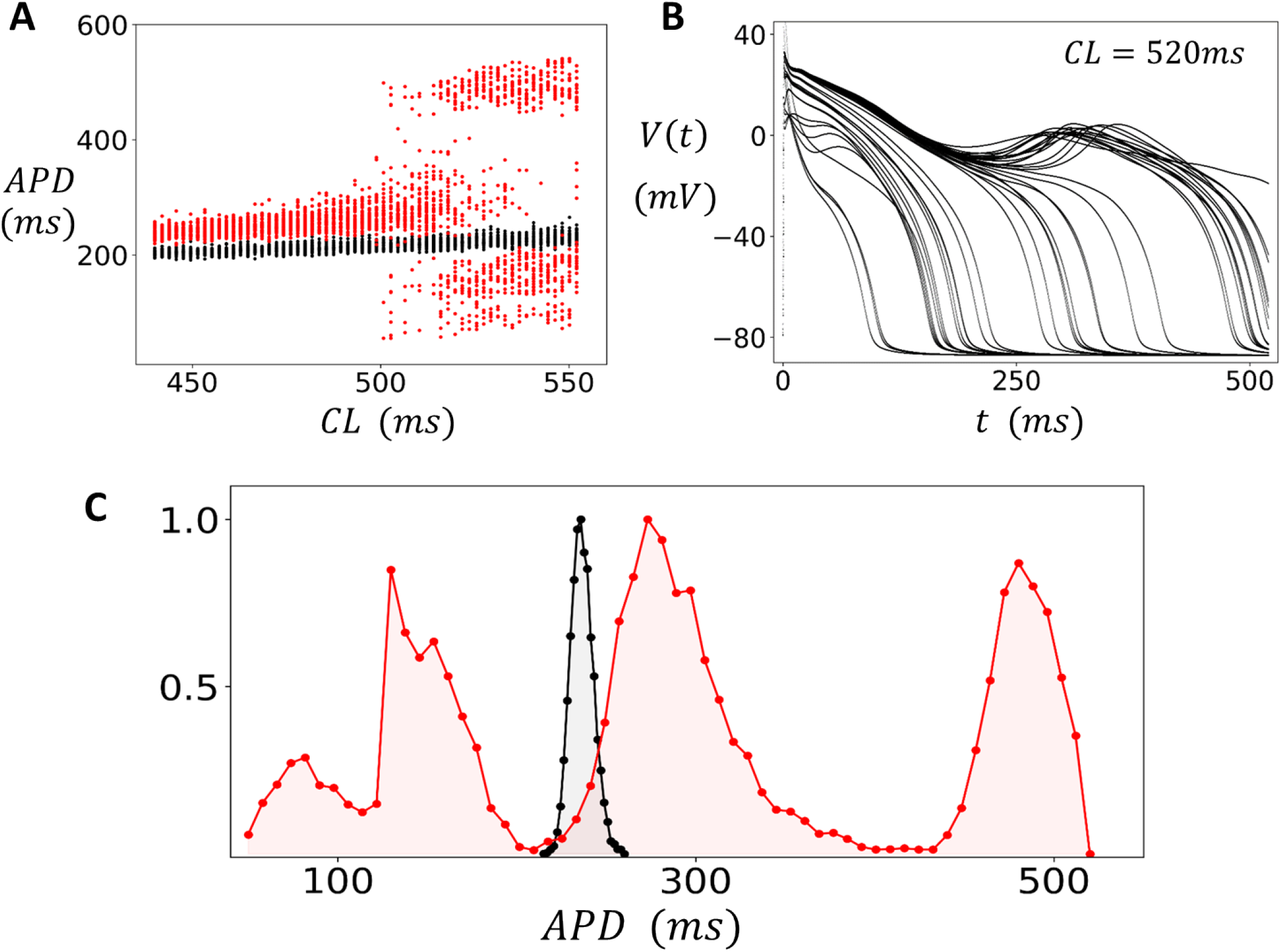
APD variability in the presence of EADs. **(A)** Bifurcation diagram using a dynamic pacing protocol. The cell is paced at a *CL* for 200 beats and the APD is plotted for the last 50 beats. Upon completion of the 200 beats the *CL* is increased incrementally and the pacing protocol is repeated. In this manner the *CL* is swept from 440*ms* to 552*ms*. The APD is measured as the time between the AP upstroke and when the voltage crosses *V*_*c*_ = ―40*mV* during repolarization. Black points denote normal conditions (*A*_*Ca*_ = 0.15), and the red points correspond to reduced Ca-induced inactivation (*A*_*Ca*_ = 0.09). The voltage time course for the last 50 beats when the cell is paced in the EAD regime (*CL* = 520*ms*). Distribution of APD measured from 15000 paced beats at *CL* = 520*ms*. The black filled circles represent the distribution under normal conditions, while the red circles denote EAD conditions. The distributions are normalized so that the maximum value is 1.

### Statistical fluctuations of the APD in 2D tissue

In cardiac tissue, neighboring cells are coupled by gap junctions, allowing voltage to spread from cell-to-cell. Consequently, the AP measured in cardiac tissue represents the spatial average of the cell population in that vicinity. The number of cells involved depends on the strength of gap junction coupling and the local arrangement of cells. In this section we apply our computational model to determine how electrical coupling modifies the distribution of the APD in the regime with and without EADs. To model tissue dynamics we solve the cable equation (Eq. 9) in a 2D tissue of size *N* × *N* cells. As a starting point we first analyze how cell coupling modifies the fluctuations of the APD in the absence of EADs. Figure 4A shows the distribution of the APD for tissue paced uniformly at *CL* = 420 for 5000 beats. In this regime EADs do not occur and the APD distribution is shown for tissue sizes of *N* = 1 (black), *N* = 5 (red), and *N* = 10 (blue). As expected, we find that electrotonic coupling substantially reduces the beat-to-beat variations in APD, and this effect becomes more pronounced as the tissue size *N* is increased. To quantify this effect more systematically, we have computed the standard deviation *σ* of the APD from the data plotted in Figure 4A, and in Figure 4B we have plotted *σ* as a function of tissue dimension *N*. This result demonstrates that the standard deviation of the APD decreases monotonically as tissue size is increased. In the regime where EADs are observed for an isolated cell (*CL* = 520*ms*) we find that the APD distribution is much broader and with a spread that depends on tissue size (Figure 4C). To explore this effect, we have also plotted the bifurcation diagram of the system (Figure 4D) where the tissue is paced to steady state for a range of *CL*. In this case we observe that in the EAD regime the APD is broadly distributed around two or three peaks with a spread that decreases with increasing tissue size.

**Figure 4.**
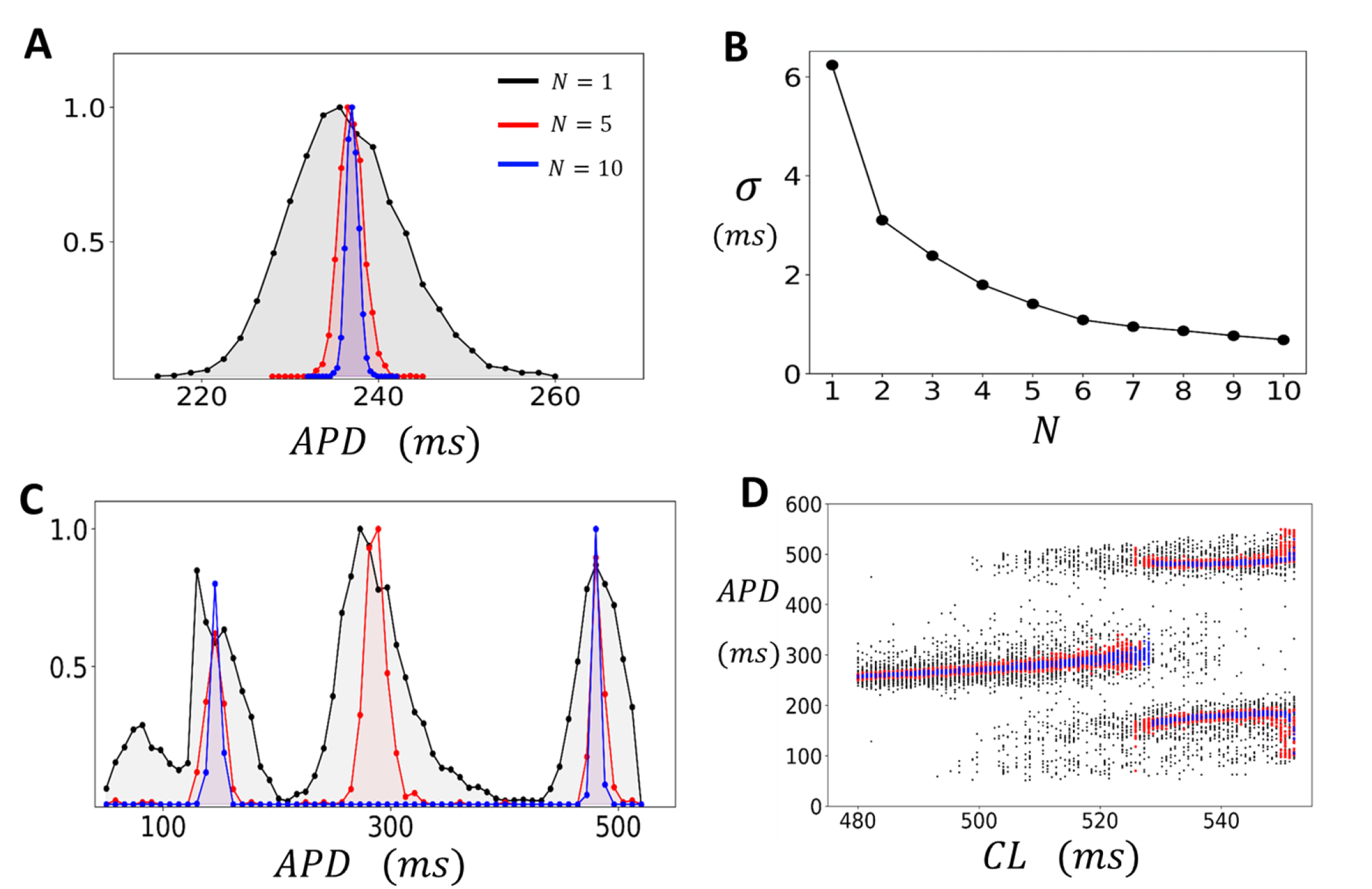
APD fluctuations in 2D tissue. **(A)** Steady state APD distribution for a 2D tissue of size *N* × *N*. The histogram is constructed by pacing the tissue at *CL* = 420*ms* for 5000 beats and recording the APD after an initial transient of 50 beats. The tissue dimension is indicated on the inset. Distributions are normalized so that the maximum is at 1. **(B)** Standard deviation of the APD (*σ*) as a function of tissue dimension *N*. **(C)** Steady state APD distribution when the tissue is paced in the EAD regime at a *CL* = 520*ms* for 5000 beats **(D)** Bifurcation diagram. The tissue is paced to steady state (200 beats) and the APD is plotted for the last 50 beats. The black, red, and blue points correspond to tissue sizes of *N* = 1,5 and 10 respectively.

### Large tissue sizes are governed by the deterministic limit of the stochastic APD model

To understand the distribution of the APD in cardiac tissue we will consider the deterministic limit of our stochastic Ca model. Our motivation is that in cardiac tissue the voltage is averaged over a population of cells so that Ca sparks are effectively recruited from a large number of J clusters. Thus, we can take the *N*_*b*_→∞ limit of equations (2-4), the number of Ca sparks in a population of cells is governed by a deterministic equation given by

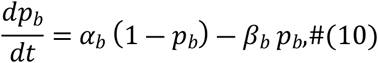

where *p*_*b*_ = *n*_*b*_/*N*_*b*_ is the fraction of J clusters with sparks. To compare the deterministic model predictions with our tissue simulations we first pace different tissue sizes to steady state and measure the average APD (⟨*APD*⟩) and standard deviation (*σ*). In Figure 5A we plot the ⟨*APD*⟩ (black circles) and *σ*, represented as an error bar around the average, when the cell is paced at *CL* = 420*ms*. On the same graph we plot the steady state APD predicted by the deterministic model (red line). Indeed, we find that as the size of the system is made larger the average APD of the stochastic model in tissue converges to the prediction of the deterministic limit. To explore the regime where the system exhibits EADs, in Figure 5B we plot the APD after the deterministic system has been paced to steady state. Specifically, the cell is paced at a given *CL* for 200 beats, and the APD of the last 50 beats is recorded. The *CL* is then increased by a small increment, and the protocol is repeated. This procedure is repeated for *CLs* in the range *CL*_0_ = 475*ms* to *CL*_1_ = 550*ms*. Once the *CL*_1_ pacing is completed the protocol is repeated, and *CL* is decreased from *CL*_1_ back to *CL*_*o*_ (red filled circles). Indeed, in this simulation we find a transition to alternans (black filled circles) when the CL is increased from *CL*_*o*_→*CL*_1_. However, when the protocol is repeated in the reverse direction *CL*_1_→*CL*_*o*_ we find that the alternans regime continues for a broader range of *CLs*. This result indicates that the steady state APD exhibits a regime of bistability. On the same plot we show the APD for the last 50 beats when the stochastic single cell model (blue dots) is paced to steady state. Here, we observe that the beat-to-beat fluctuations of APD in the single cell case are broadly scattered around the predictions of the deterministic model. Thus, at a finite system size *N*, the stochasticity from subcellular fluctuations is superimposed on the nonlinear dynamics dictated by the deterministic model. In Figure 5C we plot the voltage time course for the single cell stochastic model (black lines), and the deterministic model (red line), when the cell is paced at *CL* = 530*ms*. Indeed, we find that the single cell displays large beat to beat fluctuations in the APD, while the deterministic model exhibits APD alternans where EADs occur on alternate beats.

**Figure 5.**
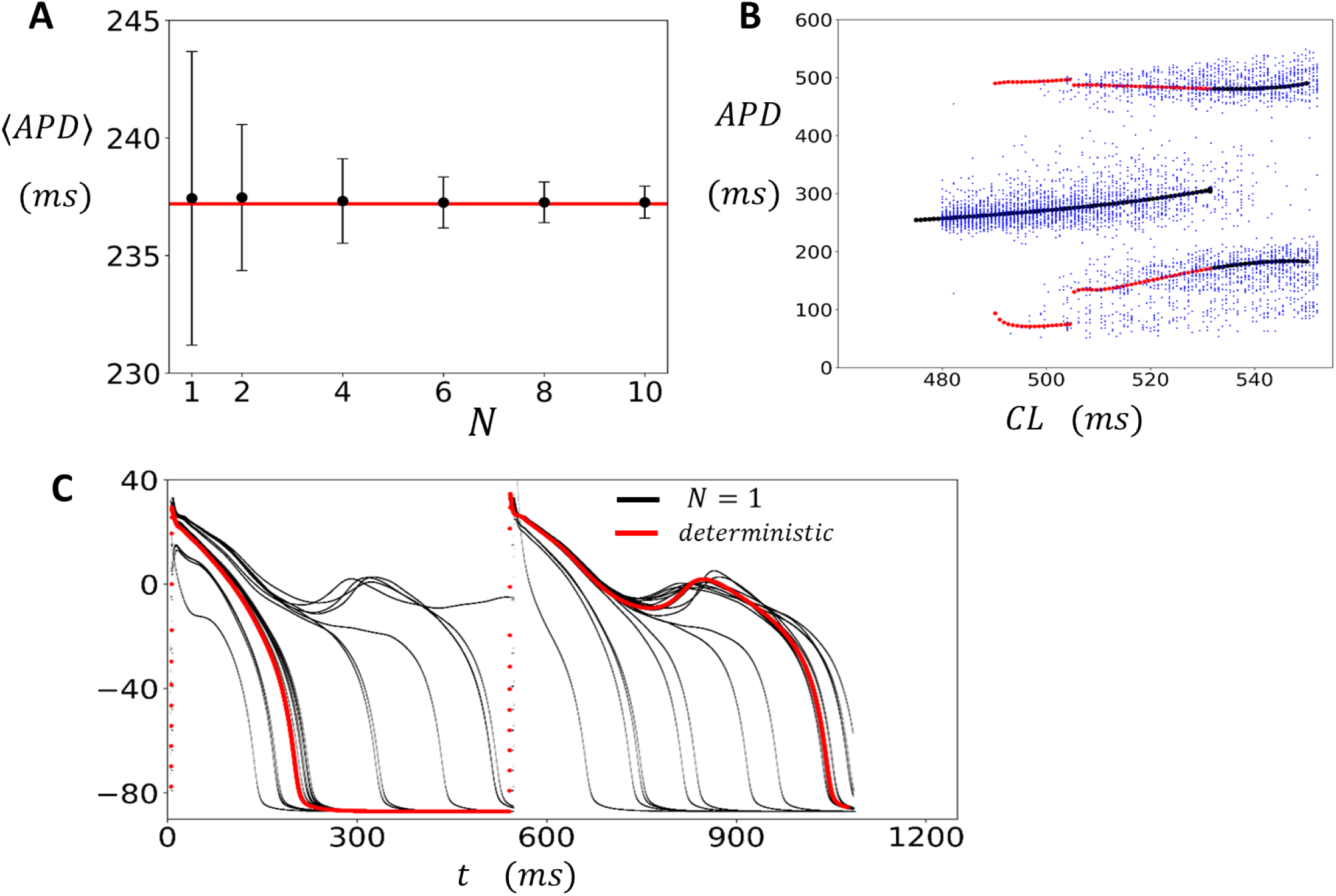
Deterministic limit of the stochastic model. **(A)** Plot of ⟨*APD*⟩ as a function of tissue size *N* (black filed circles). The standard deviation *σ* is plotted as an error bar. The tissue is paced for 5000 beats at *CL* = 420*ms*. The horizontal red line corresponds to the deterministic limit paced to steady state (50 beats). **(B)** Bifurcation diagram for the single cell stochastic model (blue filled circles) along with the predictions of the deterministic model (red and black filled circles). The deterministic bifurcation diagram is computed using a dynamic pacing protocol where the *CL* is increased gradually from *CL*_*o*_ = 475*ms* to *C L*_1_ = 550*ms* (black points). At each *CL* the cell is paced for 200 beats and the APD for the last 50 beats are plotted. Once *CL*_1_ is reached the *CL* is then decreased back to *CL*_*o*_ (red filled circles). **(C)** The steady state voltage plotted for the last 50 beats, at *CL* = 535*ms*, for the stochastic (black line) and the deterministic (red line) model.

### APD fluctuations in uniformly paced cardiac tissue

The findings from the previous section indicate that cardiac tissue exhibits beat-to-beat variations in APD due to a combination of subcellular noise and the nonlinear response of ion currents. This suggests that paced cardiac tissue will display spatial variations in the APD that is caused by these factors. To analyze these spatial patterns, we simulated a strip of cardiac tissue where all cells were paced periodically. Specifically, we paced a 3 × 100 array of coupled cells at a cycle length *CL*_1_ until steady state was reached (50 beats). We then changed the cycle length to *CL*_2_ and continued pacing at this new cycle length for 300 beats. Once a steady state was reached at *CL*_2_, we evaluated the spatial distribution of the APD on the tissue strip. In Figure 6A we plot the *APD* as a function of beat number *n* for a cell that is at the center of a tissue that is paced at *CL*_1_ = 400*ms* and *CL*_2_ = 420*ms*. In Figure 6B we plot the APD as a function of the number of cells *l*_*x*_ along the cable, after 200 (black line) and 201 (red line) beats of pacing at *CL*_2_. Here, we see that there are small spatial variations of the APD across the tissue, which are due to the underlying stochasticity of the system. In Figure 6C we show the *APD* vs *n* when the tissue is paced in the bistable regime (*CL*_2_ = 520*ms*). In this case we find that the APD exhibits large intermittent fluctuations where the APD can vary from 100*ms* to 500*ms*. In Figure 6D we show the spatial distribution of APD along the cable at beats 200 and 201. This result demonstrates that in the bistable regime the APD distribution is dynamic and can vary substantially along the cable from one beat to the next. This spatial variation is driven by fluctuation induced transitions between the two stable regimes. In Figure 6E and 6F we have paced the cable in the alternans regime where *CL* = 530*ms*. In this case the APD in tissue alternates from beat-to-beat (Figure 6E) and exhibits a stable pattern of spatially discordant alternans (Figure 6F). These results demonstrate that EADs in tissue induce dynamic spatial heterogeneity of the APD.

**Figure 6:**
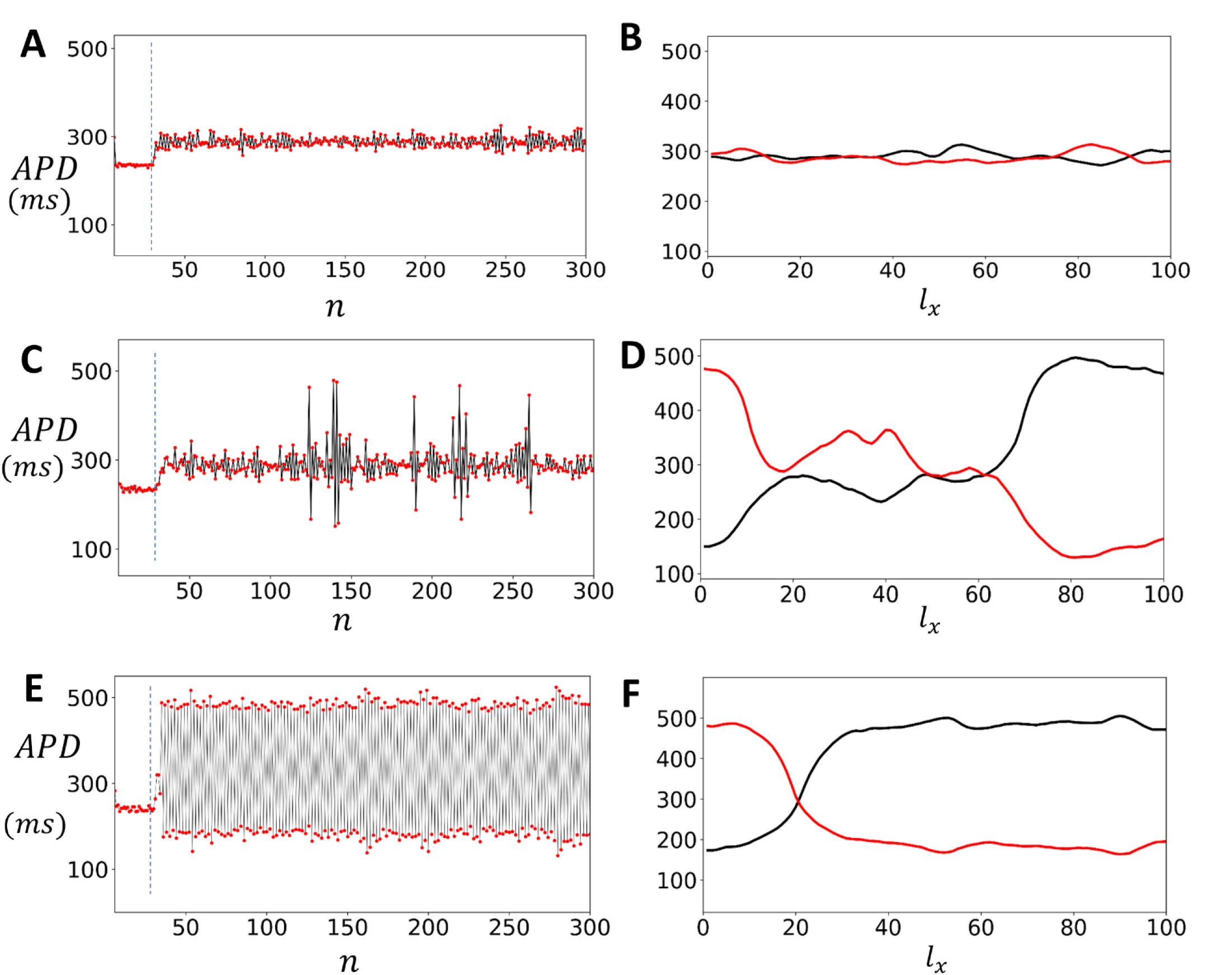
Spatial patterns and fluctuations in APD in cardiac tissue with EAD. **(A)** APD as a function of beat number *n* for a cell at the center of a 3 × 100 tissue array, paced at *CL*_1_ = 400*ms* and *CL*_2_ = 420*ms*. **(B)** APD as a function of the number of cells *l*_*x*_ along the cable after 200 (black line) and 201 (red line) beats at *CL*_2_. **(C)** APD versus *n* when the tissue is paced in the bistable regime (*CL*_2_ = 520*ms*), displaying large intermittent fluctuations. **(D)** Spatial distribution of APD along the cable at beats 200 and 201 in the bistable regime. **(E)** APD vs *n* when the system is paced in the alternans regime (*CL* = 530*ms*). **(F)** Spatial distribution of APD at beats 200 and 201 showing stable spatially discordant alternans.

### Subcellular stochasticity is highly arrhythmogenic in the EAD regime

In this section, we analyze the effect of EADs on wave propagation in 2D cardiac tissue. Figure 7 shows simulations of a 100 × 100 tissue, paced along a 10-cell-wide strip on the left edge. Initially, we pace the tissue for 20 beats at a cycle length (CL) of 420ms, where the cell model does not exhibit EADs. We then switch to a CL of 530 ms, where EADs occur in the deterministic limit. Figure 7 includes snapshots of planar wave propagation at the beat numbers indicated. By the 40th beat, the AP wave back begins to develop pronounced heterogeneities as EADs form in the tissue. These heterogeneities increase until, on the 43rd beat, the AP wavefront breaks and forms a reentrant circuit. This wave break is robust and occurs reliably across multiple simulation runs. In the discussion section, we will analyze the mechanism for wave break and show that it is a direct consequence of the spatial heterogeneity caused by EADs in cardiac tissue.

**Figure 7:**
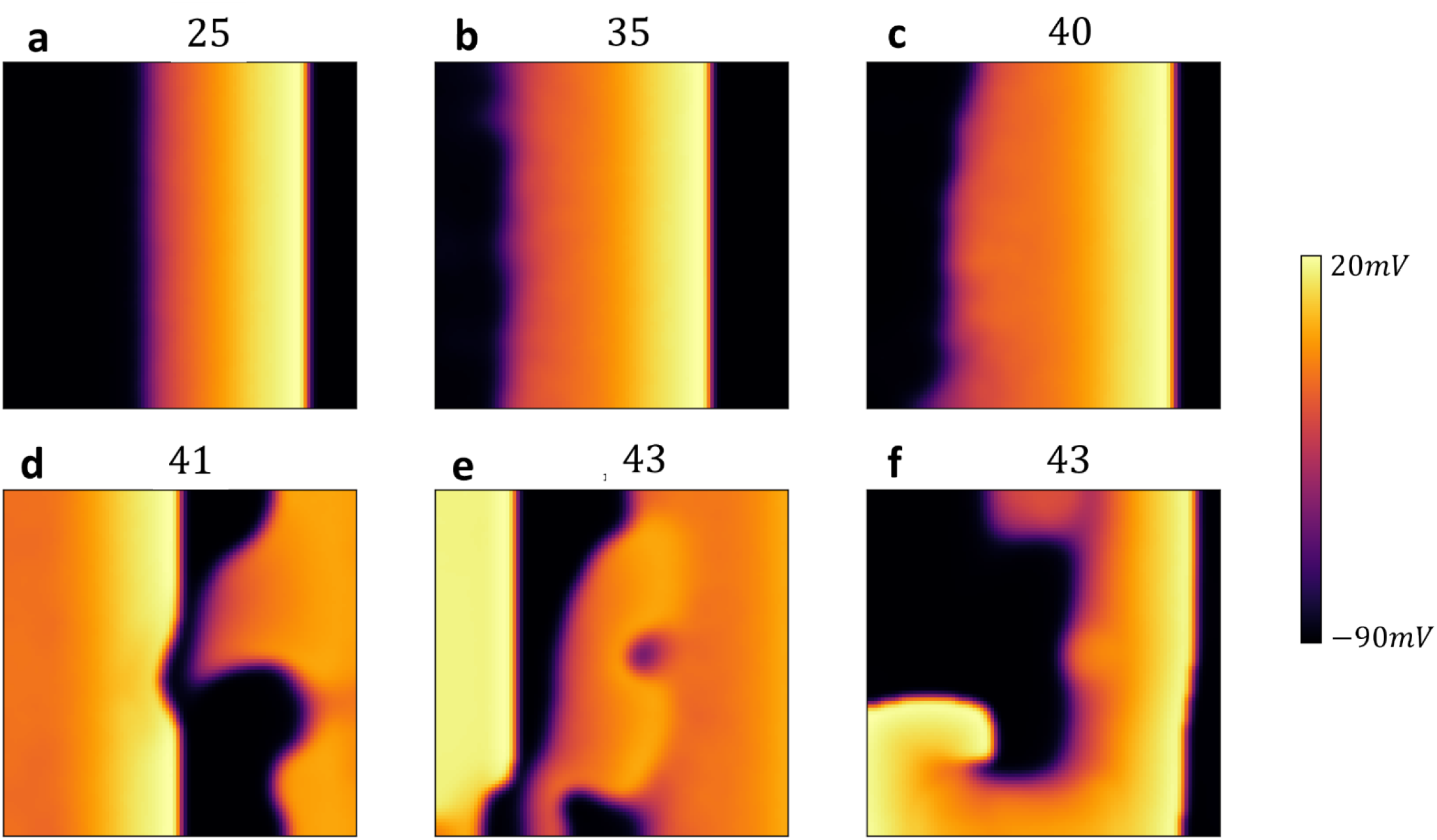
Wave Propagation in 2D Cardiac Tissue with EADs. Snapshots of planar wave propagation in a 100 × 100 cardiac tissue, paced along a 10-cell-wide strip on the left edge. The tissue is initially paced for 20 beats at *CL* = 420*ms*, where the cell model does not exhibit alternans. Subsequently, the CL is switched to530*ms*, where EADs occur in the deterministic limit. The snapshots show the development of pronounced heterogeneities in the action potential (AP) wave back by the 40th beat, leading to wavefront break and reentrant circuit formation by the 43rd beat. Snapshot **f** is 400ms after snapshot **e** during the 43rd beat.

## Discussion

### Statistics of EADs in isolated cells and tissue

In this study, we developed a computational model to describe the formation of EADs resulting from reduced calcium-induced inactivation of the LCC. The model incorporates subcellular stochasticity by accounting for the random activation and termination of calcium sparks within a population of RyR clusters in the cell. Our approach reveals that the magnitude of APD variability during pacing is sensitive to the number of RyR clusters where calcium sparks can potentially occur—specifically, those clusters that can be triggered by openings of the LCC. Our findings suggest that, to match experimental data on APD fluctuations in guinea pig myocytes, approximately 4,000 RyR clusters are required as potential sites for calcium spark initiation. This result aligns with experimental observations indicating that cardiac myocytes contain roughly 10,000 ― 20,000 RyR clusters (32-34), with only a fraction of these clusters capable of serving as calcium spark nucleation sites(35). However, it is clear that the number of clusters that can be recruited will depend on the strength of the LCC current, which itself depends on a wide range of physiological factors such as *β*-adrenergic stimulation. Thus, while APD fluctuations are dynamic and will have many sources, our study suggests that Ca spark recruitment is likely a significant contributor.

When we reduced the extent of calcium-induced inactivation by decreasing the calcium-dependent component, we observed that EADs occurred at slower pacing rates. At these rates, the APD fluctuated by over 400*ms* from beat-to-beat as EADs amplified the subcellular stochasticity. This outcome is expected, since many studies have demonstrated that EADs arise from a nonlinear instability during the AP plateau, which makes the repolarization dynamics highly sensitive to subcellular fluctuations(13,17). Our results are also consistent with experimental findings, which have shown that EADs are associated with pronounced APD variability. Notably, Li et al. (36) measured EADs over multiple beats in dog ventricular myocytes in heart failure, finding that when these myocytes were paced at 1Hz, the APD varied by up to ∼ 400*ms* during EADs. This observation is consistent with our finding that EADs in isolated cells significantly amplify subcellular fluctuations.

In cardiac tissue, cells are electrically coupled, so that the voltage of each cell will be the spatial average of a large population of neighboring cells. Thus, electrical coupling will reduce the beat-to-beat variability of the APD observed in an isolated cell. To analyze this effect more generally, we note that electrical coupling spreads over a length scale of 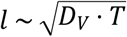, where *D*_*V*_ is the effective diffusion coefficient of voltage in tissue, and where *T* is the pacing period. Setting *T* = 500*ms* and using a standard diffusion coefficient of *D*_*V*_ ∼ 10^―4^*cm*^2^/*ms*, we find that *l* ∼ 0.2*cm*. Consequently, in cardiac tissue, the APD will represent the spatial average of the population of cells within a spherical volume of this radius, which is on the order of 10^4^ cells. Now, since the noise strength decreases as 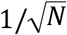, where *N* is the effective number of recruitable RyR clusters contained in that population of cells, then we have *N* ∼ 10^7^, since each cell will have roughly ∼ 10^3^ RyR clusters. Thus, in 3D tissue, the subcellular noise strength will be diminished by a factor 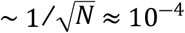, so that the voltage time course will be effectively deterministic. However, in one-dimensional (1D) strands of cardiac tissue, the situation is markedly different. Here, only about 20 cells fall within the same electrotonic length. The reduced number of cells means that stochastic effects become much more significant. As a result, fluctuations in APD due to EADs will be far more pronounced in 1D tissue than in 3D tissue. Therefore, while 3D cardiac tissue exhibits mostly deterministic behavior with minimal influence from noise, 1D cardiac tissue is heavily influenced by stochasticity, leading to substantial variability in the APD due to EADs. This finding suggests that EAD fluctuations will be considerably more pronounced in spatially restricted regions of tissue, such as Purkinje fibers(37). In these areas, we anticipate that stochastic fluctuations will induce significant spatial heterogeneities. Our simulations confirmed that in the bi-stable regime, fluctuations along a 1D cable can generate sharp heterogeneities, with different parts of the tissue shifting between distinct bi-stable states. Consequently, EADs occurring in the Purkinje fiber system may lead to pronounced spatial heterogeneities, potentially driving localized conduction block(38,39).

### Nonlinear dynamics and EADs

A crucial finding of this study is that in 2D cardiac tissue, APD fluctuates around values determined by the deterministic limit of the stochastic model. Thus, in order to understand the statistical distribution of the APD, it is necessary to characterize the beat-to-beat dynamics of the deterministic model. To achieve this, we will construct a nonlinear map model of the APD, which will identify the key nonlinear properties driving the instability. Our approach is to determine the functional dependence of the APD on the previous diastolic interval, denoted as *DI*, by computing the S1S2 restitution curve of the model. This approach has been used in previous studies to analyze period doubling bifurcations in cardiac cells(40). To construct the S1S2 restitution curve, we first pace the cell at the S1 cycle length. Once the cell has reached steady state, we vary the CL of the last paced beat, referred to as the S2 stimulus, and measure the final APD and the preceding *DI*. This procedure is repeated for different S2 durations and in Figure 8A, we plot the final APD versus *DI*. The S1S2 restitution curve for our model indicates that the APD increases gradually as a function of *DI*, until roughly *DI* ∼ 350*ms*, after which it increases abruptly. To explain the shape of the restitution curve, in the inset we show the voltage time course for the last paced beat, corresponding to the four representative points on the restitution curve. This shows that the steep increase of the APD is due to the onset of EADs which induces long APDs when *DI* approaches a critical value. Here, we note that the same shape of the APD restitution curve has been observed in previous studies where EADs are induced by different ionic mechanisms(18,41). Thus, this APD restitution shape is not specific to our model, but is expected more generally in ionic models which exhibit long APDs characteristic of EADs. To explore the effect of this nonlinearity we have fitted a cubic spline to the restitution curve (Figure 8A, black line) and used this function to construct a nonlinear map of the form

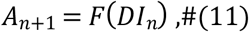

where *A*_*n*_ and *DI*_*n*_ denote the APD and DI at beat *n*, and which is iterated at fixed CL denoted as *T* = *DI*_*n*_ + *A*_*n*_. In Figure 8B, we present the bifurcation diagram for the nonlinear map, showing a transition from a stable periodic fixed point to a period doubled steady state. To generate the diagram, we iterated the map at fixed values of *T*, incrementally increasing *T* after reaching steady state, and plotted the steady-state APD (blue circles). Red circles represent data obtained by decreasing *T* from the period-doubled solution. These results indicate that the period-doubling bifurcation is bistable and that the periodic fixed point loses stability discontinuously. To analyze these features further in Figure 8C we show cobweb diagrams for *T* = 550*ms* using two initial conditions: *DI*_*o*_ = 350*ms* and *DI*_*o*_ = 450*ms*. This result indicates that the period doubled solution corresponds to EADs occuring only on alternate beats. These results are consistent with the bifurcation diagram of the full ionic model, which also exhibited a similar bistable period-doubling bifurcation. Finally, we emphasize that the bifurcation observed here is not unique to our model. A similar bifurcation has been reported in other models of EADs involving different mechanisms. For example, Liu et al. (41) investigated a Long QT syndrome model and found that the transition to EADs occurred through a discontinuous transition to EAD alternans. This suggests that the transition we describe is a generic feature of all EAD models and not confined to a specific cellular mechanism.

**Figure 8:**
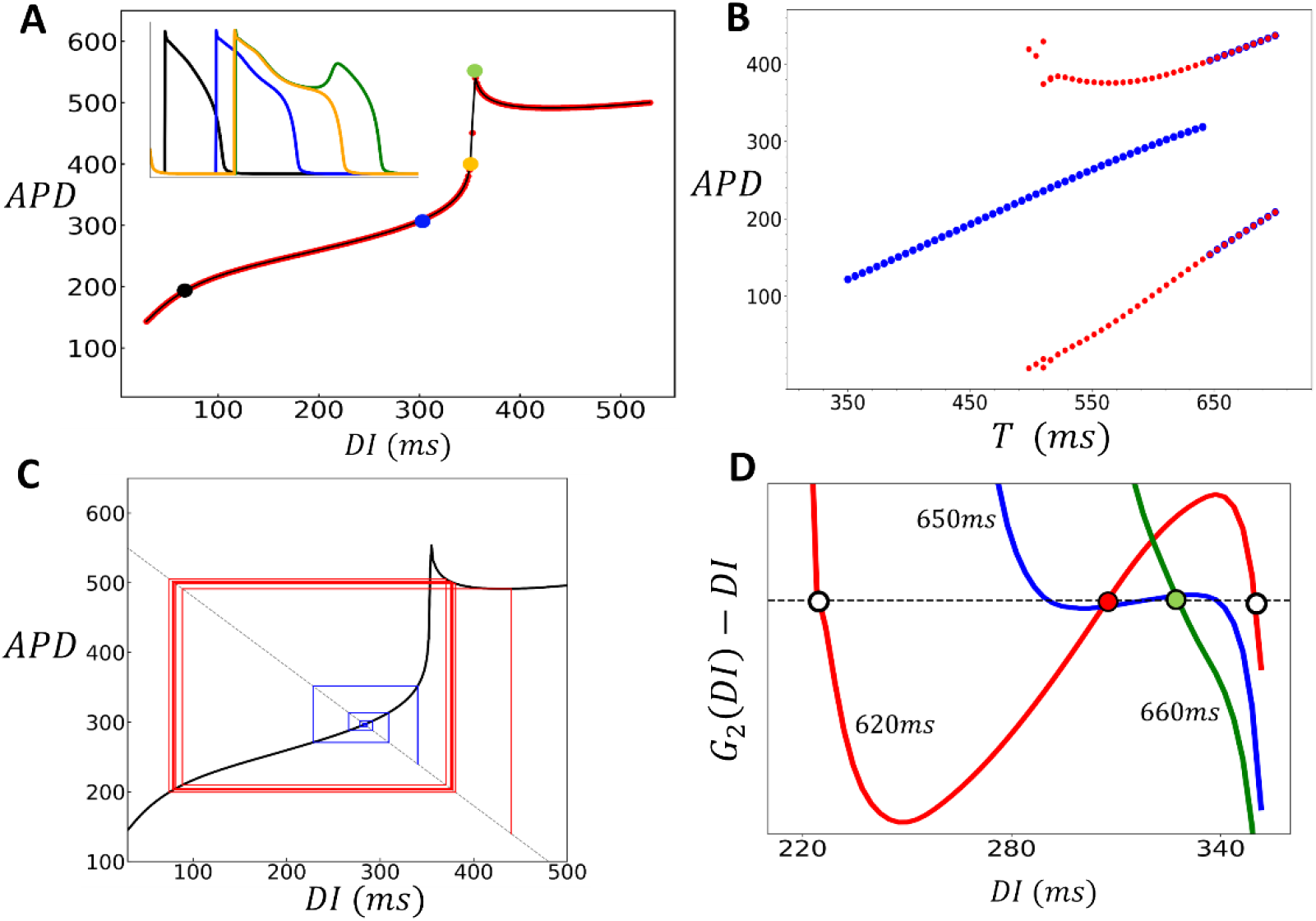
Analysis of Nonlinear Dynamics and EAD-Induced Bifurcations in Cardiac Tissue. **(A)** S1S2 restitution curve. The final APD is plotted against the previous diastolic interval (*DI*) after pacing the cell to steady state (50 beats) at *S*1 = 500*ms*. The red points are the measured APD for the S2 beat and the black line is a cubic splines fit. The inset shows voltage for the last paced beat at the indicated points on the restitution curve. The black, blue, yellow, and green plots correspond to the colored points on the restitution curve. **(B)** Period-doubling bifurcation diagram. The presence of EADs leads to a period-doubling bifurcation, resulting in an alternating behavior where EADs occur only on alternate beats. Blue points are computed by sweeping the period from low to high, and the red points are computed by sweeping the period from high to low. **(C)** A cob-web diagram demonstrating bistability of the nonlinear map where the stable fixed point (blue) coexists with an alternating behavior (red). The pacing period is fixed at *T* = 550*ms*. **(D)** Plot of the function *G*_2_(*DI*) ―*DI*, where *G*_2_(*DI*) is the second iteration of the S1S2 restituion map. Intersections with the dashed line indicate fixed points of the second iteration of the nonlinear map. At *T* = 620*ms* (red line) the system posseses two unstable fixed points (white circles) and one stable fixed point (red circle). At *T* = 660*ms* (green line) the system only possesses one unstable fixed point (green circle). This transition occurs occurs near *T* ≈ 650*ms* where the stable fixed point loses its stability via a subcritical pitch fork bifurcation (blue line). Note that this plot only shows the range of *DI* near the stable fixed point, as the system possesses two other stable fixed points corresponding to the bistable state.

Our nonlinear map analysis uncovers several important characteristics of the transition to EAD alternans in the deterministic limit. The first important observation of the transition, in both the nonlinear map and the full ionic model, is that it is discontinuous. To explain this feature, we apply standard methods to analyze how the periodic fixed point of the map bifurcates into the period doubled solution. Our approach is to rewrite Eq. 11 as the single variable map *DI*_*n*+1_ = *G*(*DI*_*n*_), where *G*(*DI*) = *T* ― *F*(*DI*). To analyze the period doubled solutions we consider the second iteration of the map, *G*_2_(*DI*) = *G*(*G*(*DI*)), so that a period doubled solution satisfies the condition *DI* = *G*_2_(*DI*). In Figure 8D we study the solutions of *G*_2_(*DI*) ―*DI* = 0 as the pacing period *T* is varied across the transition. The red line shows the function *G*_2_(*DI*) ―*DI* for the case *T* = 620*ms* which is less than the onset of the period doubling bifurcation at *T*_*c*_ ∼ 650*ms*. Here, we see that there are two unstable solutions (white circles), and one stable solutions (red circle). However, above the transition, where *T* = 660*ms* (green line), we find that there is only a single stable solution (green circle). Also, we plot the function near onset at *T* = 650*ms* (blue line) to see that the two unstable fixed points vanish at the instant where the stable fixed point loses stability. This transition is known as a subcritical pitchfork bifurcation, which is characterized by a discontinuous jump to alternans. This type of bifurcation typically occurs when the system exhibits an approximate reflection symmetry around the stable fixed point. In our system, this symmetry is roughly satisfied due to the shape of the restitution curve near the onset of EADs. This behavior is in sharp contrast to the transcritical bifurcation seen in the standard APD restitution instability to alternans (1), where APD alternans develop gradually without a discontinuous jump.

Our nonlinear analysis of EADs in cardiac tissue reveals several findings worth noting. First, in paced cardiac tissue, EADs appear as alternans, occurring only on every other beat. This occurs because cardiac tissue can be effectively modeled by the deterministic limit of our stochastic model, where EADs arise following a period-doubling bifurcation. Importantly, this alternating behavior is observed only at the tissue level. In isolated cells subcellular noise introduces significant variability in EAD occurrence and a consistent alternans response will not be observed. Second, this bifurcation is discontinuous so that even an infinitesimal change in pacing rate leads to a large change in the APD. This feature causes EADs in tissue to occur abruptly since they emerge after a discontinuous transition. Finally, the transition to EAD alternans is bistable, so that the alternating EAD alternans can coexist with a stable periodic state. This result has important physiological implications. In particular, the onset of EADs will be highly sensitive to pacing rate and tissue heterogeneities. Thus, when tissue becomes prone to EADs they will likely emerge in a highly heterogenous fashion. Also, since the transition is bistable different parts of tissue can exhibit either a periodic or alternating response. These features ensure that when cardiac tissue is paced near the EAD threshold will exhibit steep APD gradients which are highly arrhythmogenic.

### EADs promote conduction block in paced cardiac tissue

Our pacing simulations in 2D cardiac tissue (Figure 7) revealed that when EADs occur, a planar excitation becomes highly unstable, leading to conduction block and re-entry. To better understand this phenomenon, we computed the S1S2 restitution curve in a strip of cardiac tissue, using the same S1S2 pacing protocol as described in the previous section. To generate the restitution curve, we paced the left end of the tissue at a fixed cycle length (CL) and measured the APD at the distal end. We then applied the S1S2 protocol and measured the APD as a function of the preceding *DI* for the final paced beat. The resulting restitution curve is shown in Figure 9A. This curve is similar to the single cell restitution curve with the difference that when *DI* falls below approximately 25*ms*, the AP fails to propagate, resulting in conduction block. The range where propagation fails is highlighted by the green shaded box in the figure. The presence of a conduction block regime implies that an S2 beat can fail to propagate if delivered bellow a fixed threshold. In Figure 9A, this is illustrated using cobweb diagrams for two different initial conditions, shown in red and blue. When the S2 is applied with an intermediate *DI* (blue line), the system converges to a stable fixed point. However, when the *DI* is larger (red line), the electrical excitation fails to propagate, resulting in conduction block. This occurs due to the distinctive shape of the APD restitution curve, which leads to large APDs when EADs are present. As a result, a long APD prevents the tissue from adequately recovering, causing conduction block on the following beat. To further investigate this effect across a range of initial conditions, we used a cubic spline fit to the APD restitution curve and analyzed how different initial conditions evolve over many beats. In Figure 9B we have computed the phase diagram of the system. The y-axis represents the initial diastolic interval *DI*_0_, while the x-axis represents the pacing period (*T*). For each initial condition, the nonlinear map was iterated 100 times to determine the system’s long-term behavior. In the resulting phase diagram, blue regions indicate a stable period-1 solution, red regions represent a stable period-2 solution, and black regions denote the occurrence of conduction block. This result demonstrates that APD propagation in tissue with EADs is extremely sensitive to small changes in the pacing period. Therefore, in paced cardiac tissue— where the timing of an excitation wavefront is expected to vary—the system becomes highly prone to wavebreak. This sensitivity explains our observations (Figure 7) of robust wavebreak during pacing of 2D cardiac tissue.

**Figure 9:**
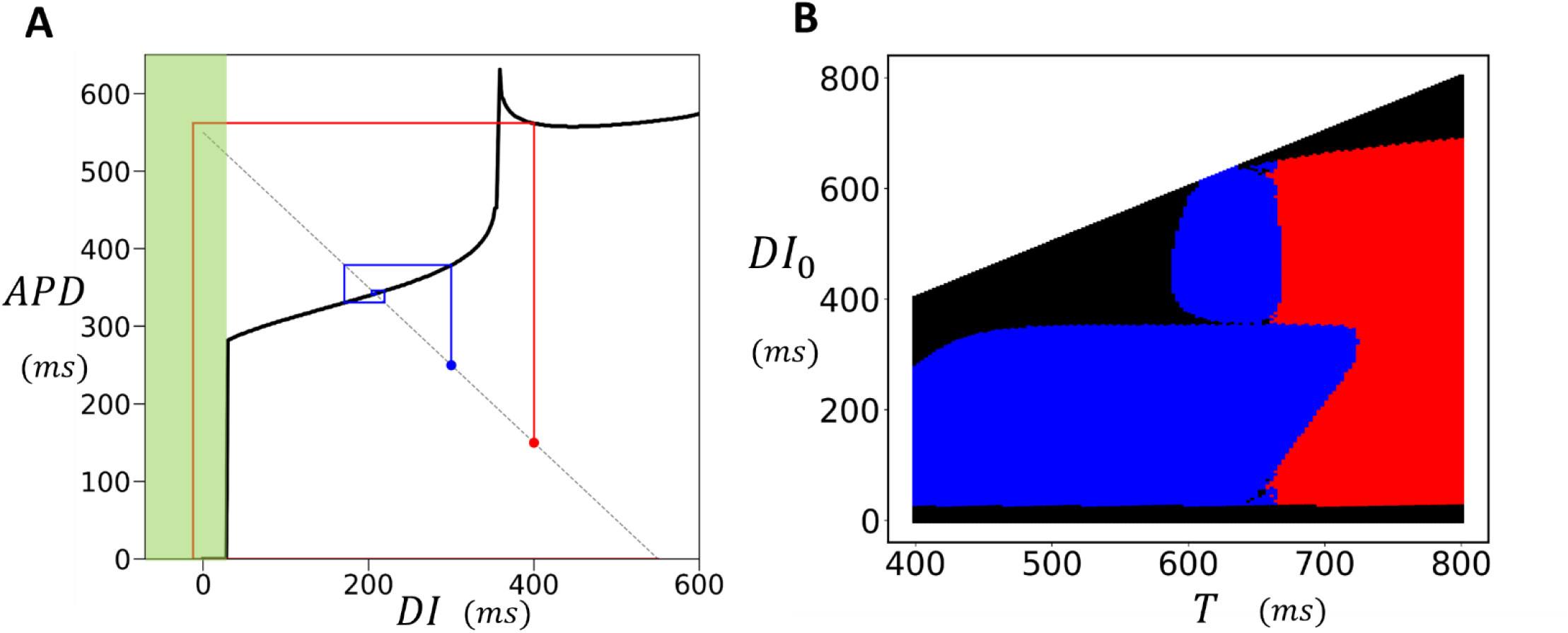
Conduction block in Cardiac Tissue. **(A)** S1S2 restitution curve for a tissue strip 60 × 3 cells. All cells on a strip of size 10 × 3 are stimulated simulatenously using an S1S2 protocol and the APD is measured at the distal end at cell (60, 2). Black line denotes a cubic splines fit to the S1S2 restitution curve. Conduction block occurs for S2 intervals shorter than approximately *DI* ≈ 25 *ms* (green shaded region). Cobweb diagrams are drawn using a cubic spline fit to the restitution curve for initial conditions *DI*_0_ = 300*ms* (blue line) and *D*_0_ = 400*ms* (red line). Here, the red line indicates a sequence leading to conduction block. **(B)** Phase diagram of the system using the cubic splines fit to the S1S2 restitution curve. The y-axis represents the initial diastolic interval *DI*_0_, while the x-axis represents the pacing period *T*. For each initial condition, the nonlinear map was iterated 100 times. In the resulting phase diagram, blue regions indicate a stable period-1 solution, red regions indicate a stable period-2 solution, and black regions represent conduction block i.e. the system reaches *DI* < 25*ms* at some point during the 100 iteration trajectory.

Our analysis suggests that EADs in tissue occur via alternans which promotes conduction block and reentry. However, several studies have suggested that EADs can induce arrhythmias via propagating ectopic beats(42,43). In this scenario an EAD in tissue propagates into nearby refractory tissue, and that excitation proceeds to form reentry. Though this scenario is possible, the conditions for successful propagation of an EAD in cardiac tissue are quite stringent. For instance, Deo et al. (44) analyzed ectopic excitations resulting from EADs in the Purkinje system and showed that these excitations can lead to reentry, but only under precise conditions. Specifically, they identified strict requirements on the timing of the excitation and the electrophysiological state of the substrate for reentry to occur. In contrast, our study of paced cardiac tissue revealed that reentry occurred robustly following the onset of EADs without requiring ectopic excitations. Instead, it was the heterogeneity introduced by EAD alternans that led to conduction block, which then facilitated reentry. Therefore, our findings suggest that the more likely mechanism of EAD-induced arrhythmogenesis is through EAD alternans and conduction block, rather than ectopic excitation. However, a comprehensive quantitative analysis is still needed to fully determine which mechanism—ectopic excitation or conduction block driven by alternans—dominates in the induction of arrhythmias under different physiological conditions.

## Conclusion

In conclusion, our study provides a comprehensive study of EADs and their effects at both the cellular and tissue levels. We find that EADs in isolated cells are highly variable due to subcellular stochastic processes, leading to significant beat-to-beat variability in the APD. However, in electrically coupled cardiac tissue, these fluctuations are largely dampened, and the dynamics can be described by a deterministic model. Our main finding is that EADs in tissue manifest as alternans, where EADs occur on every other beat. This transition to EAD alternans is discontinuous and leads to pronounced spatial heterogeneities which are strongly arrhythmogenic. These findings provide critical insights into the mechanisms by which cellular stochasticity can cause tissue-level arrhythmogenic events, emphasizing the importance of both nonlinear dynamics and electrical coupling in arrhythmia onset and propagation.

## Author contributions

All authors contributed equally to the conception, design, execution, and interpretation of the research.

## Acknowledgement

This work was supported by the National Institute of General Medical Sciences (Award Number: 1R16GM153647-01) and the National Science Foundation (Award Number: 2320846) to YS. FF was supported by NIH (Award Number: 2R01HL143450-05A1). We thank these agencies for their generous support.

## REFERENCES

1. Weiss, J. N., Z. Qu, P.-S. Chen, S.-F. Lin, H. S. Karagueuzian, H. Hayashi, A. Garfinkel, and A. Karma. 2005. The dynamics of cardiac fibrillation. Circulation. 112(8):1232–1240.

2. Haissaguerre, M., E. Vigmond, B. Stuyvers, M. Hocini, and O. Bernus. 2016. Ventricular arrhythmias and the His–Purkinje system. Nature Reviews Cardiology. 13(3):155–166.

3. January, C. T., J. M. Riddle, and J. J. Salata. 1988. A model for early afterdepolarizations: induction with the Ca2+ channel agonist Bay K 8644. Circulation research. 62(3):563–571.

4. January, C. T., and A. Moscucci. 1992. Cellular Mechanisms of Early Afterdepolarizations a. Annals of the New York Academy of Sciences. 644(1):23–32.

5. Rosen, M. R., J. P. Moak, and B. Damiano. 1984. The clinical relevance of afterdepolarizations. Annals of the New York Academy of Sciences. 427(1):84–93.

6. Weiss, J. N., A. Garfinkel, H. S. Karagueuzian, P.-S. Chen, and Z. Qu. 2010. Early afterdepolarizations and cardiac arrhythmias. Heart rhythm. 7(12):1891–1899.

7. Wit, A. L. 2018. Afterdepolarizations and triggered activity as a mechanism for clinical arrhythmias. Pacing and Clinical Electrophysiology. 41(8):883–896.

8. Roden, D. 1993. Early after-depolarizations and torsade de pointes: implications for the control of cardiac arrhythmias by prolonging repolarization. European heart journal. 14(suppl_H):56–61.

9. Choi, B. R., F. Burton, and G. Salama. 2002. Cytosolic Ca2+ triggers early afterdepolarizations and Torsade de Pointes in rabbit hearts with type 2 long QT syndrome. The Journal of physiology. 543(2):615–631.

10. Roden, D. M. 2016. Predicting drug-induced QT prolongation and torsades de pointes. The Journal of physiology. 594(9):2459–2468.

11. Fowler, E. D., N. Wang, M. Hezzell, G. Chanoit, J. C. Hancox, and M. B. Cannell. 2020. Arrhythmogenic late Ca2+ sparks in failing heart cells and their control by action potential configuration. Proceedings of the National Academy of Sciences. 117(5):2687–2692.

12. Zeng, J., and Y. Rudy. 1995. Early afterdepolarizations in cardiac myocytes: mechanism and rate dependence. Biophysical journal. 68(3):949–964.

13. Tran, D. X., D. Sato, A. Yochelis, J. N. Weiss, A. Garfinkel, and Z. Qu. 2009. Bifurcation and chaos in a model of cardiac early afterdepolarizations. Physical review letters. 102(25):258103.

14. Kimrey, J., T. Vo, and R. Bertram. 2020. Canard analysis reveals why a large Ca2+ window current promotes early afterdepolarizations in cardiac myocytes. PloS Computational Biology. 16(11):e1008341.

15. Wang, R., Z. Qu, and X. Huang. 2024. Dissecting the roles of calcium cycling and its coupling with voltage in the genesis of early afterdepolarizations in cardiac myocyte models. PLOS Computational Biology. 20(2):e1011930.

16. Huang, X., Z. Song, and Z. Qu. 2018. Determinants of early afterdepolarization properties in ventricular myocyte models. PLoS computational biology. 14(11):e1006382.

17. Sato, D., L.-H. Xie, A. A. Sovari, D. X. Tran, N. Morita, F. Xie, H. Karagueuzian, A. Garfinkel, J. N. Weiss, and Z. Qu. 2009. Synchronization of chaotic early afterdepolarizations in the genesis of cardiac arrhythmias. Proceedings of the National Academy of Sciences. 106(9):2983–2988.

18. Sato, D., L.-H. Xie, T. P. Nguyen, J. N. Weiss, and Z. Qu. 2010. Irregularly appearing early afterdepolarizations in cardiac myocytes: random fluctuations or dynamical chaos? Biophysical journal. 99(3):765–773.

19. Zaniboni, M., A. E. Pollard, L. Yang, and K. W. Spitzer. 2000. Beat-to-beat repolarization variability in ventricular myocytes and its suppression by electrical coupling. American Journal of Physiology-Heart and Circulatory Physiology. 278(3):H677–H687.

20. Shiferaw, Y., G. L. Aistrup, and J. A. Wasserstrom. 2018. Synchronization of triggered waves in atrial tissue. Biophysical journal. 115(6):1130–1141.

21. Shiferaw, Y., G. L. Aistrup, and J. A. Wasserstrom. 2017. Mechanism for triggered waves in atrial myocytes. Biophysical journal. 113(3):656–670.

22. Pásek, M., F. Brette, A. Nelson, C. Pearce, A. Qaiser, G. Christé, and C. H. Orchard. 2008. Quantification of t-tubule area and protein distribution in rat cardiac ventricular myocytes. Progress in biophysics and molecular biology. 96(1-3):244–257.

23. Ibrahim, M., J. Gorelik, M. H. Yacoub, and C. M. Terracciano. 2011. The structure and function of cardiac t-tubules in health and disease. Proceedings of the Royal Society B: Biological Sciences. 278(1719):2714–2723.

24. Bootman, M. D., I. Smyrnias, R. Thul, S. Coombes, and H. L. Roderick. 2011. Atrial cardiomyocyte calcium signalling. Biochimica et Biophysica Acta (BBA)-Molecular Cell Research. 1813(5):922–934.

25. Chen-Izu, Y., S. L. McCulle, C. W. Ward, C. Soeller, B. M. Allen, C. Rabang, M. B. Cannell, C. W. Balke, and L. T. Izu. 2006. Three-dimensional distribution of ryanodine receptor clusters in cardiac myocytes. Biophysical journal. 91(1):1–13.

26. Dibb, K. M., J. D. Clarke, M. A. Horn, M. A. Richards, H. K. Graham, D. A. Eisner, and A. W. Trafford. 2009. Characterization of an extensive transverse tubular network in sheep atrial myocytes and its depletion in heart failure. Circulation: Heart Failure. 2(5):482–489.

27. Mahajan, A., Y. Shiferaw, D. Sato, A. Baher, R. Olcese, L.-H. Xie, M.-J. Yang, P.-S. Chen, J. G. Restrepo, and A. Karma. 2008. A rabbit ventricular action potential model replicating cardiac dynamics at rapid heart rates. Biophysical journal. 94(2):392–410.

28. Qi, X., Y.-H. Yeh, D. Chartier, L. Xiao, Y. Tsuji, B. J. Brundel, I. Kodama, and S. Nattel. 2009. The calcium/calmodulin/kinase system and arrhythmogenic afterdepolarizations in bradycardia-related acquired long-QT syndrome. Circulation: Arrhythmia and Electrophysiology. 2(3):295–304.

29. Jin, Q., J. L. Greenstein, and R. L. Winslow. 2022. Estimating the probability of early afterdepolarization and predicting arrhythmic risk associated with long QT syndrome type 1 mutations. Biophysical Journal. 121(3):89a.

30. Clusin, W. T. 2003. Calcium and cardiac arrhythmias: DADs, EADs, and alternans. Critical reviews in clinical laboratory sciences. 40(3):337–375.

31. Qu, Z., and A. Garfinkel. 1999. An advanced algorithm for solving partial differential equation in cardiac conduction. IEEE transactions on biomedical engineering. 46(9):1166–1168.

32. Soeller, C., D. Crossman, R. Gilbert, and M. B. Cannell. 2007. Analysis of ryanodine receptor clusters in rat and human cardiac myocytes. Proceedings of the National Academy of Sciences. 104(38):14958–14963.

33. Baddeley, D., I. Jayasinghe, L. Lam, S. Rossberger, M. B. Cannell, and C. Soeller. 2009. Optical single-channel resolution imaging of the ryanodine receptor distribution in rat cardiac myocytes. Proceedings of the National Academy of Sciences. 106(52):22275–22280.

34. Shen, X., J. van den Brink, Y. Hou, D. Colli, C. Le, T. R. Kolstad, N. MacQuaide, C. R. Carlson, P. M. Kekenes-Huskey, and A. G. Edwards. 2019. 3D dSTORM imaging reveals novel detail of ryanodine receptor localization in rat cardiac myocytes. The Journal of Physiology. 597(2):399–418.

35. Cheng, H., and W. Lederer. 2008. Calcium sparks. Physiological reviews. 88(4):1491–1545.

36. Li, G.-R., C.-P. Lau, A. Ducharme, J.-C. Tardif, and S. Nattel. 2002. Transmural action potential and ionic current remodeling in ventricles of failing canine hearts. American Journal of Physiology-Heart and Circulatory Physiology. 283(3):H1031–H1041.

37. Nogami, A., Y. Komatsu, A. K. Talib, W. Phanthawimol, Q. J. Naeemah, T. Haruna, and I. Morishima. 2023. Purkinje-related ventricular tachycardia and ventricular fibrillation: solved and unsolved questions. Clinical Electrophysiology. 9(10):2172–2196.

38. Boyden, P. A., W. Dun, and R. B. Robinson. 2016. Cardiac Purkinje fibers and arrhythmias; the GK moe award lecture 2015. Heart rhythm. 13(5):1172–1181.

39. Boyden, P. A. 1996. Cellular electrophysiologic basis of cardiac arrhythmias. The American journal of cardiology. 78(4):4–11.

40. Kalb, S. S., H. M. Dobrovolny, E. G. Tolkacheva, S. F. Idriss, W. Krassowska, and D. J. Gauthier. 2004. The restitution portrait: A new method for investigating rate-dependent restitution. Journal of cardiovascular electrophysiology. 15(6):698–709.

41. Liu, W., T. Y. Kim, X. Huang, M. B. Liu, G. Koren, B. R. Choi, and Z. Qu. 2018. Mechanisms linking T-wave alternans to spontaneous initiation of ventricular arrhythmias in rabbit models of long QT syndrome. The Journal of physiology. 596(8):1341–1355.

42. Li, Z.-Y., C. Maldonado, C. Zee-Cheng, S. Hiromasa, and J. Kupersmith. 1992. Purkinje fibre-papillary muscle interaction in the genesis of triggered activity in a guinea pig model. Cardiovascular research. 26(5):543–548.

43. El-Sherif, N., R. H. Zeiler, W. Craelius, W. B. Gough, and R. Henkin. 1988. QTU prolongation and polymorphic ventricular tachyarrhythmias due to bradycardia-dependent early afterdepolarizations. Afterdepolarizations and ventricular arrhythmias. Circulation research. 63(2):286–305.

44. Deo, M., P. M. Boyle, A. M. Kim, and E. J. Vigmond. 2010. Arrhythmogenesis by single ectopic beats originating in the Purkinje system. American Journal of Physiology-Heart and Circulatory Physiology. 299(4):H1002–H1011.

